# Predictive inference alterations in psychosis proneness are context-dependent

**DOI:** 10.1101/2025.05.17.654666

**Authors:** Luca Tarasi, Vincenzo Romei

## Abstract

**Background:** Perception depends on the dynamic integration of sensory inputs and prior expectations, shaped by recent experience. While disruptions in this inferential process have been proposed as core features of psychosis, whether such alterations can be observed in subclinical psychosis-like experiences (PLEs), and how they manifest across different forms of perceptual inference remains unclear.

**Methods:** Two independent samples of healthy adults participated in the study: 80 completed a visual detection task, and 62 performed a motion discrimination task. Data were analysed using mixed-effects models with a continuous PLE measure and supplemented by categorical comparisons between the lowest- and highest-PLEs terciles.

**Results:** PLEs predicted reduced sensitivity to sensory evidence and increased reliance on probabilistic cues across both tasks. Critically, history biases diverged across contexts. In the detection task, a repulsive bias from the previous stimulus was observed in the overall sample, but this effect was attenuated in high-PLEs. In contrast, in the discrimination task, high-PLEs exhibited a repulsive bias away from their previous choice, reversing the typical repetition tendency seen in low-PLEs.

**Conclusions:** Our results suggest that psychosis-like traits alter how prior expectations interact with sensory input and recent perceptual history. High-PLEs showed reduced influence of recent sensory input in detection task, consistent with altered low-level adaptation. In contrast, in the discrimination task, they showed a shift away from previous choices, suggesting disruption of decision-level integration of past choices. This points to a task-dependent instability in the use of past information, reflecting a broader inferential vulnerability linked to the psychosis spectrum.

## Introduction

Perception is not a passive readout of sensory input, but an active inferential process whereby the brain integrates sensory evidence with prior knowledge to generate probabilistic estimates of the external world. This view, formalized in predictive processing theories, holds that perceptual experience arises from the interplay between bottom-up signals and top-down predictions, shaped by the minimization of prediction error (1,2). Crucially, when the balance between sensory evidence and prior expectations becomes dysregulated, perception itself can become distorted —leading individuals to perceive expected stimuli in the absence of reliable evidence, or to misinterpret ambiguous inputs in line with internal predictions (3,4). Growing evidence suggests that such computational imbalances lie at the core of psychosis-spectrum phenomena, spanning from attenuated symptoms to full-blown schizophrenia (5,6). For example, hallucinations have been proposed to reflect an overreliance on prior beliefs, which dominate perception even in the absence of reliable sensory input (3,7). Delusions, in turn, may emerge when low-level sensory priors are imprecise, allowing spurious signals to gain undue significance. This uncertainty may trigger a compensatory shift at higher hierarchical levels, where cognitive priors become overly precise, constraining interpretation in rigid, belief-consistent ways (8,9). These aberrations in perceptual inference reflect a fundamental miscalibration in the weighting of internal predictions versus external input.

Importantly, similar computational deviations have been documented in the general population. Psychosis-like experiences (PLEs)—unusual perceptual or belief-related phenomena occurring in the absence of clinical disorder—are increasingly recognized as meaningful subclinical expressions of psychosis vulnerability (10,11). Recent findings indicate that individuals with elevated PLEs exhibit reduced sensory precision and an exaggerated reliance on prior expectations—paralleling the inferential signatures observed in schizophrenia (12–15). These results support a dimensional view of psychosis, in which disruptions to predictive coding mechanisms unfold along a continuum and may be detectable well before the emergence of clinical symptoms.

In addition to the integration of priors and sensory evidence, perceptual inference also relies critically on recent experience. Humans display robust sequential biases, whereby current choices are influenced by preceding stimuli and responses (16–19). These serial dependencies are increasingly viewed as expressions of predictive inference, reflecting the brain’s tendency to exploit temporal regularities to optimize perception under uncertainty. Although sequential biases typically enhance perceptual stability, they appear disrupted in psychosis, with altered history effects reported in both clinical and high-risk populations (20–22), but see (23). A key open question is whether these alterations reflect a stable, trait-like vulnerability marker, or rather a context-sensitive maladaptation—whereby perceptual history is used inconsistently depending on task demands.

To date, no study has systematically investigated how psychosis proneness shapes the integration of sensory evidence, prior expectations, and choice history across tasks that differ in their hierarchical and computational demands. Critically, while reduced serial dependence has often been interpreted as a general marker of impaired predictive processing in psychosis, such accounts may overlook the possibility that these biases also reflect a maladaptive sensitivity to task structure—leading to unstable or context-inappropriate inference strategies depending on the hierarchical nature of the decision.

Here, we directly address this gap by combining two well-characterized paradigms that probe distinct forms of perceptual uncertainty. In the first, a visual detection task, participants judge the presence or absence of near-threshold stimuli. In the second, a motion discrimination task, they resolve directional uncertainty using dynamic dot patterns that require continuous evidence accumulation (24). Although both tasks were titrated to yield comparable accuracy, they differ fundamentally in their computational demands and the level of processing they engage. Specifically, the detection task places greater demands on early sensory encoding, where perceptual history may influence subsequent trial via low-level adaptation mechanisms. In contrast, the discrimination task draws more heavily on internal decision dynamics, making it more sensitive to how previous choices influence current behaviour. This distinction provides a framework to test whether psychosis-like traits differentially affect the integration of past information across the sensory–decisional axis—disrupting stimulus-driven biases when adaptation is key, and altering choice-related biases in tasks that rely more heavily on decision-level processing.

Using generalized linear mixed models, we quantified the influence of current sensory input, prior cues, previous stimuli, and previous choices on perceptual decisions and, critically, how these factors interact with individual differences in psychosis proneness. By integrating both continuous and tercile-based analyses of PLEs, our approach captures dimensional and categorical signatures of altered predictive coding across the psychosis spectrum.

We show that elevated PLEs are associated with reduced sensitivity to sensory evidence and increased reliance on prior expectations across tasks, consistent with predictive coding accounts. Crucially, sequential biases revealed a striking task-dependent dissociation: in the detection task, high-PLE individuals showed a marked attenuation of the typical repulsive bias from the previous stimulus, whereas in the discrimination task, they exhibited a repulsive bias relative to their previous choice—reversing the attractive bias seen in low-PLE participants. These results indicate that psychosis-prone individuals do not exhibit a uniform reduction in serial dependence, but rather a disruption in the integration of past information across different inferential contexts. Such fragmented use of sensory and decisional history may reflect early disruptions in the hierarchical architecture supporting predictive inference.

## Methods

We conducted two behavioural experiments to examine how prior expectations and sequential dependencies influence perceptual decisions as a function of psychosis proneness. A total of 142 healthy adults participated (Experiment 1: *n* = 80; Experiment 2: *n* = 62). Participants with mean accuracy ±2 standard deviations from the group mean were excluded to minimize the influence of performance outliers on model estimation (Exp. 1: *n* = 5; Exp. 2: *n* = 2).

### Experiment 1: Detection Task

Participants performed a probabilistic detection task designed to assess how perceptual decisions are shaped by varying levels of sensory expectation (25,26) (Figure 1A). The task comprised two phases. In the initial phase, an adaptive psychophysical procedure was used to determine each participant’s contrast threshold for detecting grey circular targets. This titration yielded ∼70% accuracy across a balanced mix of target-present and catch trials (27). Stimuli were presented in a dimly lit room using MATLAB (R2016b) and the Psychophysics Toolbox. In the main experiment, each trial began with a visual probabilistic cue indicating the likelihood that a target would appear. When present, the target consisted of a grey circle embedded within a checkerboard background, displayed in the lower-left quadrant of the screen for 60 ms. Each trial began with a cue presented for a fixed duration of 1 s, followed by a variable interval of 1.2– 1.5, after which the checkerboard appeared either with or without the target. Participants were instructed to respond using their right hand, pressing the “K” key if they detected the target, or “M” if they perceived its absence. No time pressure was imposed. After each response, a black screen was presented for a randomly varied interval ranging between 1.9 and 2.4 seconds. The probabilistic cue took the form of a vertically split rectangle, shaded red on the bottom and blue on the top. The proportion of red visually encoded the likelihood of target occurrence. There were three cue conditions: high probability, low probability and a neutral condition, which provided no predictive information. Importantly, the actual frequency of target appearances across conditions matched the probabilities conveyed by the cues, and participants were explicitly informed about this correspondence.

**Fig. 1.**
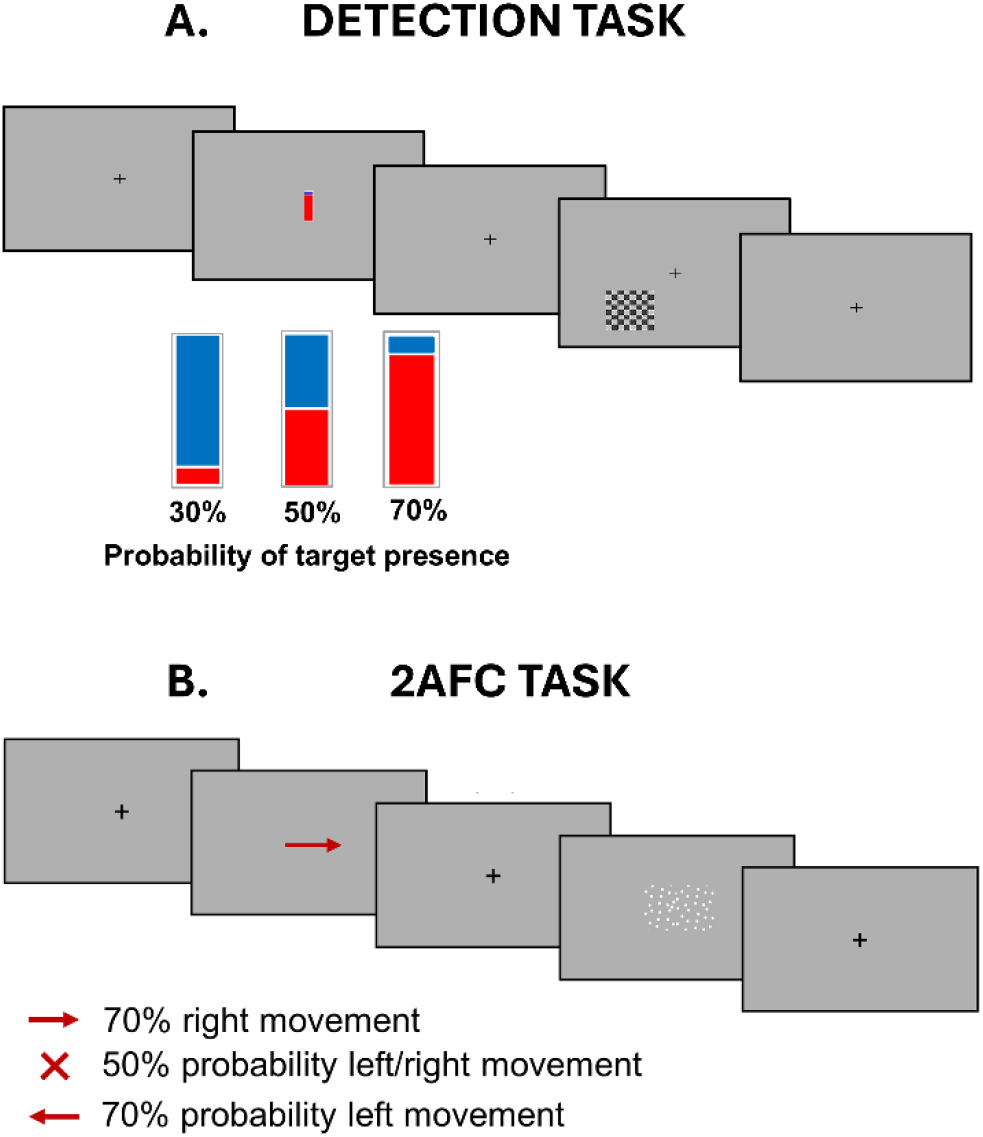
Probabilistic Detection task and 2AFC Random Dot Motion Task **A**. Each trial began with a central cue—a vertically split bar (red bottom, blue top)—displayed for 1 s. The bar fill level indicated the probability of target presence: high (70%), low (30%), or neutral (50%). After a variable delay (1.2–1.5 s), a checkerboard with or without grey circles appeared for 0.06 s, followed by a black screen (1.9–2.4 s). **B**. Each trial began with a probabilistic cue indicating the likelihood of leftward or rightward motion. After a 1 s delay, a centrally presented dot motion stimulus appeared for 0.4 s. Cues were either informative (arrows indicating 70% probability) or uninformative (a cross indicating 50%). In both tasks, cue validity matched actual target probability, and participants were informed accordingly. Stimulus contrast (checkerboard) and motion coherence (dots) were titrated to achieve ∼70% accuracy.

### Experiment 2: Random Dot Motion discrimination task

Participants performed a motion discrimination task using dynamic random-dot stimuli composed of 400 white dots displayed within a central square region (28); Figure 1B). Stimuli moved at a fixed velocity of 4.5°/s and were presented on a monitor positioned 70 cm from the seated participants in a dim environment, utilizing MATLAB (v2016b) and Psychophysics Toolbox. The stimuli varied in coherence—the proportion of dots moving uniformly either to the left or right—across several difficulty levels (0%, 3%, 6%, 9%, 15%, 21%, 30%, and 60%). At 0% coherence, dots moved randomly, whereas at higher coherence levels, an increasing proportion moved consistently in one direction, thus manipulating task difficulty. An individualized coherence threshold, corresponding to 70% accuracy, was determined beforehand for each participant via a titration phase employing the constant stimuli method and fitting a psychometric function with the *psignifit* toolbox (29). In each experimental trial, participants indicated the perceived dot motion direction (left or right) by pressing designated keys (F5 for leftward, F12 for rightward) using their index finger. Before the main task, participants completed a brief demonstration followed by a training phase to familiarize themselves with task requirements. The main experiment consisted of four blocks of 120 trials each, during which directional expectations were manipulated by probabilistic cues presented 1 second before stimulus onset. Cues consisted of left- or right-pointing arrows (informative cues, indicating increased likelihood of the corresponding target location) or a central cross (uninformative cue, indicating equal likelihood). Each trial began with a fixation period lasting 3.5 s, followed by the probabilistic cue and the subsequent stimulus presented at the pre-determined coherence level for 400 ms. Participants were explicitly informed of cue validity to ensure the correct interpretation of the probability information provided.

### Questionnaires

In both experiments, participants completed the Schizotypal Personality Questionnaire (SPQ) (30), a widely used self-report measure of schizotypal traits in non-clinical populations. The SPQ comprises subscales targeting cognitive-perceptual, interpersonal, and disorganized features associated with the schizophrenia spectrum. For this study, we focused on a composite index of psychosis-like experiences (PLEs), computed by summing scores from three subscales: Magical Thinking, Ideas of Reference, and Unusual Perceptual Experiences. Mean PLE scores were 5.91 ± 5.33 (SD) in Experiment 1 and 2.92 ± 3.44 in Experiment 2. These were selected for their strong theoretical and empirical relevance to anomalous beliefs and sensory distortions thought to index subclinical expressions of psychosis. This PLE index served as an individual-differences variable to examine how variability in psychosis-proneness modulates perceptual decision-making and the integration of prior expectations.

### Statistical Analyses

Behavioral responses were analyzed using a generalized linear mixed-effects model (GLMM) with a binomial distribution and logit link function, implemented in MATLAB R2021a via the fitglme function. The dependent variable was the participant’s binary choice (present vs. absent in Experiment 1; leftward vs. rightward in Experiment 2). The logistic model was defined as follows:

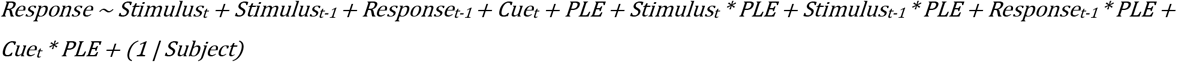

History-dependent effects were modeled by including lagged regressors for the previous stimulus and response (Stimulus_t−1_ and Response_t−1_). The model included trial-level predictors—current stimulus (Stimulus_t_), previous stimulus (Stimulus_t−1_), previous response (Response_t−1_), and cue probability (Cue)—as well as a subject-level predictor indexing psychosis-like experiences (PLE). Interaction terms were defined a priori based on our hypotheses and following the approach of Eckert et al. (20). Specifically, we tested interactions between PLE and: (i) previous response (choice history), (ii) previous stimulus (stimulus history), (iii) cue probability (expectations), and (iv) current stimulus (sensory evidence). To preserve the temporal structure of the data, trials were sorted by subject and trial number; the first trial for each participant was excluded due to missing lagged information. A random intercept was included to account for inter-individual variability.

### Group-based Tercile analysis

To complement the dimensional analysis based on continuous PLE scores, we conducted a follow-up analysis in which participants were stratified into terciles according to their composite PLE scores. Individuals falling within the lowest and highest terciles were assigned to Low and High PLE groups, respectively. A generalized linear mixed-effects model (GLMM) was then fitted separately for each task, using the same structure as the main model but replacing the continuous PLE predictor with a group factor (PLEgroup: Low = 0, High = 1). Fixed effects included current stimulus (Stimulus_t_), previous response (Response_t−1_), previous stimulus (Stimulus_t−1_), cue probability (Cue_t_), and their interactions with PLEgroup. Random intercepts were specified for each participant. This approach enabled us to test the robustness of the effects observed in the main model—specifically, how psychosis proneness modulates sensory processing, sequential biases, and expectation use—when comparing individuals at opposite ends of the schizotypy continuum.

## Results

### Behavioural performance

Participants successfully completed two perceptual decision-making tasks designed to dissociate the influence of bottom-up sensory evidence, top-down expectations, and trial history, and to examine how these factors are modulated by psychosis-like experiences (PLEs). Performance was calibrated using individualized titration procedures, yielding near-threshold accuracy in both experiments (Experiment 1: mean = 74%, SD = 7.1%; Experiment 2: mean = 71.2%, SD = 5.2%). To quantify the contribution of sensory, sequential, and expectation-related factors, we fitted generalized linear mixed-effects models (GLMMs) to each dataset. Fixed effects included current stimulus, previous stimulus, previous response, and cue probability, as well as their interactions with PLE scores; random intercepts were specified for each participant. Trials were sorted by subject and trial number, with the first trial excluded to preserve lag structure. Variance Inflation Factors (VIFs) were all below 1.4 (Exp. 1: max VIF = 1.32; Exp. 2: max VIF = 1.21), indicating no multicollinearity issues.

### Perceptual decisions are shaped by sensory evidence, prior cues, and sequential context

Across both experiments, participants’ decisions were primarily driven by current sensory input. In the detection task, the presence of a target significantly increased the probability of reporting “present” (β = 2.55 ± 0.042, p < .001). A similarly strong effect of stimulus direction was observed in the motion discrimination task (β = 1.75 ± 0.041, p < .001), confirming the dominant role of bottom-up evidence in guiding perceptual reports. In addition to sensory input, participants integrated probabilistic cues into their decisions. In both tasks, higher cue probability increased the likelihood of responding in the expected direction (Exp 1: β = 0.50 ± 0.02, p < .001; Exp 2: β = 0.47 ± 0.03, p < .001), consistent with flexible incorporation of prior beliefs. Responses were also influenced by trial history, although this effect varied across tasks. In the detection task, participants showed a strong tendency to repeat the previous response (β = 0.86 ± 0.05, p < .001), indicating a robust choice bias. Conversely, a significant repulsive effect of the previous stimulus was observed (β = –0.29 ± 0.05, p < .001). This means that participants’ responses are biased away from the previous trial’s stimulus. For example, if the target was present in trial_t − 1_, they were more likely to report its absence in trial_t_ (and vice versa). In contrast, while a similar repulsive effect of prior stimulus emerged in the motion task (β = –0.22 ± 0.04, p < .001), the main effect of previous response was not significant (β = 0.05 ± 0.04, p = .30), suggesting weaker sequential choice dependencies in this context.

### Dimensional psychosis-like traits modulate reliance on sensory input and prior knowledge

Critically, individual differences in psychosis proneness modulated decision-making processes in both tasks. Participants with higher PLE scores exhibited reduced reliance on sensory evidence, as reflected in a significant interaction between stimulus and PLE (Exp. 1: β = –0.047 ± 0.005, p < .001; Exp. 2: β = –0.038 ± 0.009, p < .001). This attenuation suggests that individuals higher in psychosis-proneness were less influenced by bottom-up information, even when the sensory evidence was task-relevant. Concurrently, the influence of top-down expectations was amplified in those with higher PLE scores. A significant positive interaction was found between cue probability and PLE in both experiments (Exp. 1: β = 0.011 ± 0.003, p < .001; Exp. 2: β = 0.051 ± 0.006, p < .001), indicating that prior beliefs had a stronger impact on decision outcomes in individuals with elevated PLE traits. This reciprocal modulation of top-down and bottom-up processing provides strong empirical support for an imbalance in predictive mechanisms underlying perceptual anomalies in psychosis proneness.

### The influence of choice history depends on task context

Interestingly, PLE exerted differential effects on history biases across tasks. In the detection task, PLE did not significantly modulate choice repetition (Response_t-1_ × PLE: β = 0.007 ± 0.006, p = .221), but did reduce the repulsive influence of the previous stimulus (Stimulus_t-1_ × PLE: β = 0.015 ± 0.005, p = .006), suggesting that psychosis‐prone individuals are less likely to bias away from the last trial’s sensory input. By contrast, in the motion-discrimination task, PLE scores were specifically associated with a repulsive bias away from the previous response (Response_t−1_ × PLE: β = –0.020 ± 0.001, p = .046), indicating a reduced tendency to repeat past choices and, instead, a shift toward alternating decisions. No modulatory effect of the previous stimulus was observed (Stimulus_t−1_ × PLE: β = 0.004 ± 0.010, p = .698), suggesting that, in individuals with high PLE, sequential biases selectively emerge at the decisional level rather than the sensory-adaptation level in this task. Together, these results reveal a double dissociation whereby psychosis-like traits disrupt stimulus-based adaptation in tasks relying on early sensory encoding and reshape response-based biases in tasks that depend on higher-order mechanisms. This pattern highlights a task-contingent breakdown in the temporal integration of perceptual history, pointing to a hierarchical disorganization in how past information is leveraged to support stable decision-making over time.

### Tercile-based effects of PLE on decision variables

The tercile-based analysis confirmed and extended the effects observed with continuous PLE scores. In the detection task (Experiment 1), high-PLE participants showed a markedly reduced reliance on current sensory evidence (Stimulus_t_ × PLEgroup: β = –0.631 ± 0.063, *p* < .001), indicating that individuals in the top tercile weighted bottom-up input substantially less than their low-PLE counterparts. Cue effects remained facilitatory in the high-PLE group (Cue_t_ × PLEgroup: β = 0.114 ± 0.039, *p* = .004), and there was no significant modulation of choice repetition by PLEgroup (Response_t−1_ × PLEgroup: β = 0.102 ± 0.075, *p* = .176), consistent with the continuous model. Crucially, serial repulsion from the previous stimulus was attenuated in high-PLE participants (Stimulus_t−1_ × PLEgroup: β = 0.164 ± 0.073, *p* = .024), suggesting a reduction of the observed repulsive bias in those with elevated psychosis‐like traits.

In the motion discrimination task (Experiment 2), high-PLE individuals again exhibited increased reliance on explicit cues (Cue_t_ × PLEgroup: β = 0.477 ± 0.050, *p* < .001), alongside a reduced weighting of sensory evidence (Stimulus_t_ × PLEgroup: β = –0.210 ± 0.076, *p* = .006). Notably, they also showed a significant reduction in response repetition (Response_t−1_ × PLEgroup: β = –0.338 ± 0.085, *p* < .001), indicating a bias away from their previous choice. There was a marginal interaction between previous stimulus and PLE group (Stimulus_t−1_ × PLEgroup: β = 0.155 ± 0.084, *p* = .065), indicating a trend-level modulation of stimulus history effects by psychosis proneness.

Taken together, these findings indicate that high-PLE individuals consistently down-weight sensory input and place increased reliance on probabilistic expectations across tasks. However, the expression of history-dependent biases diverges: in detection, a reduction of stimulus-based repulsion points to disrupted adaptation-like encoding, while in discrimination, altered response repetition reflects a shift in higher-order decisional strategies. These results reinforce the view that psychosis proneness disrupts the temporal integration of past information in a hierarchical and context-sensitive manner.

**Fig. 2.**
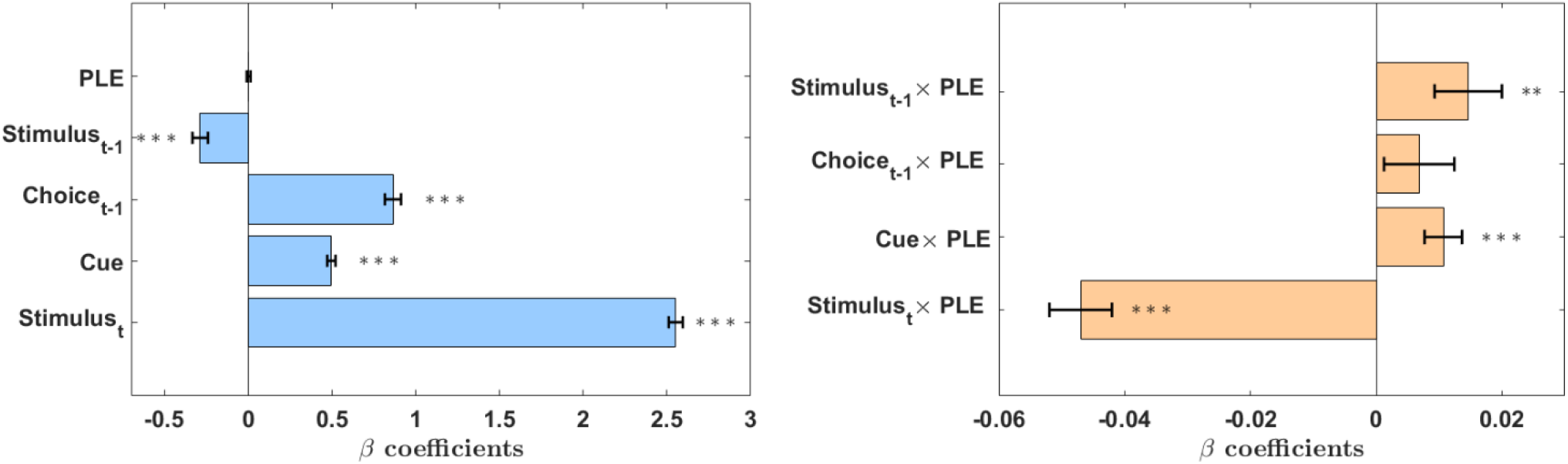
Detection task: Coefficients of the logistic choice model. In the left panel, both the current stimulus (Stimulus_t_) and cue probability (Cue_t_) exert significant positive influences on choice, while the previous stimulus (Stimulus_t−1_) shows a significant repulsive effect, and the previous choice (Choice_t−1_) has a significant positive effect, indicating a tendency to repeat the previous response. Psychosis-proneness (PLE) does not show a significant main effect. In the right panel, higher PLE scores are associated with reduced sensitivity to current sensory evidence (Stimulus_t_ × PLE), increased reliance on the cue (Cue_t_ × PLE) and diminished repulsion from the previous stimulus (Stimulus_t−1_ × PLE), while the effect of previous choice is not significantly modulated by PLE (Choice_t−1_ × PLE). Colored bars indicate β estimates and black lines represent SEM. Abbreviations: Stimulus_t_/Stimulus_t−1_ = current/previous stimulus, Choice_t_/Choice_t−1_ = current/previous choice, Cue = explicit probabilistic information, PLE = psychosis‐proneness score. Significance: ^*^p < .05;^**^p < .01; ^***^p < .001.

**Fig. 3.**
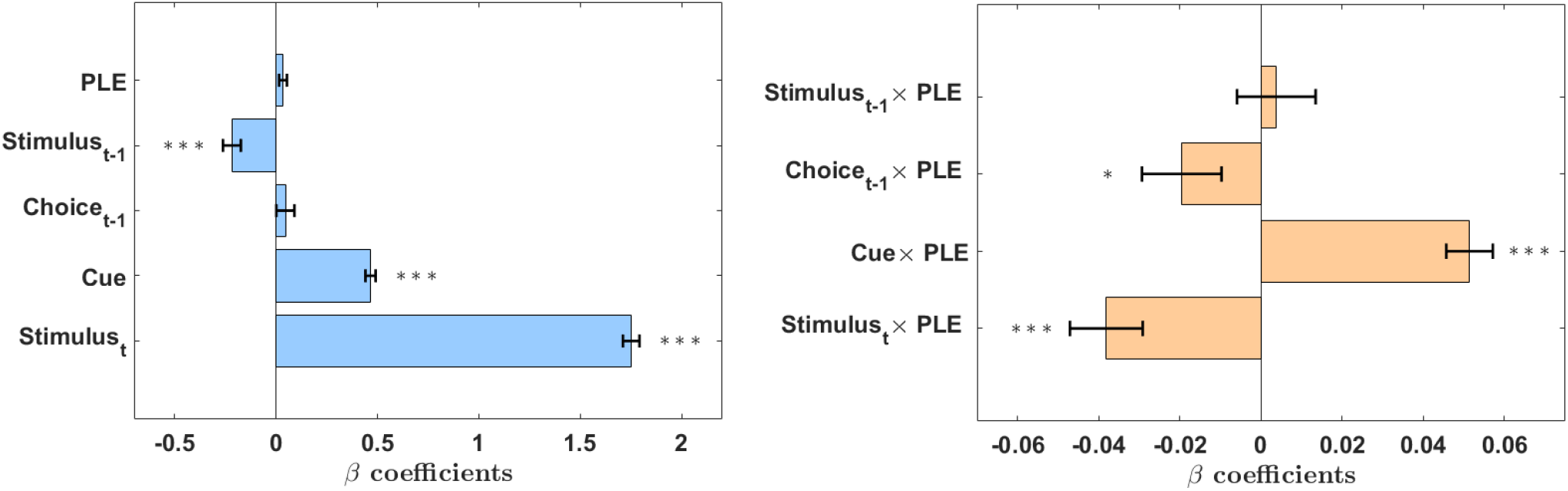
Motion discrimination task: Coefficients of the logistic choice model. In the left panel, current sensory evidence (Stimulus_t_) and cue probability (Cue_t_) both exert significant positive influences on choice, the preceding stimulus (Stimulus_t−1_) yields a negative (repulsive) effect, while neither previous choice (Choice_t−1_) nor psychosis‐proneness (PLE) shows significant main effect. In the right panel, higher PLE scores significantly reduce sensitivity to current sensory evidence (Stimulus_t_ × PLE) and significantly enhance reliance on the cue (Cue_t_ × PLE). No PLE-related modulation was observed for the effect of the previous stimulus (Stimulus_t−1_ × PLE), whereas higher PLE scores were associated with a reduced tendency to repeat previous responses—indeed, a shift toward repulsion (Choice_t−1_ × PLE). Colored bars indicate β estimates and black lines represent SEM. Abbreviations: Stimulus_t_/Stimulus_t−1_ = current/previous stimulus, Choice_t_/Choice_t−1_ = current/previous choice, Cue = explicit probabilistic information, PLE = psychosis‐proneness score. Significance: ^*^ p < .05;^**^p < .01; ^***^p < .001.

**Fig. 4.**
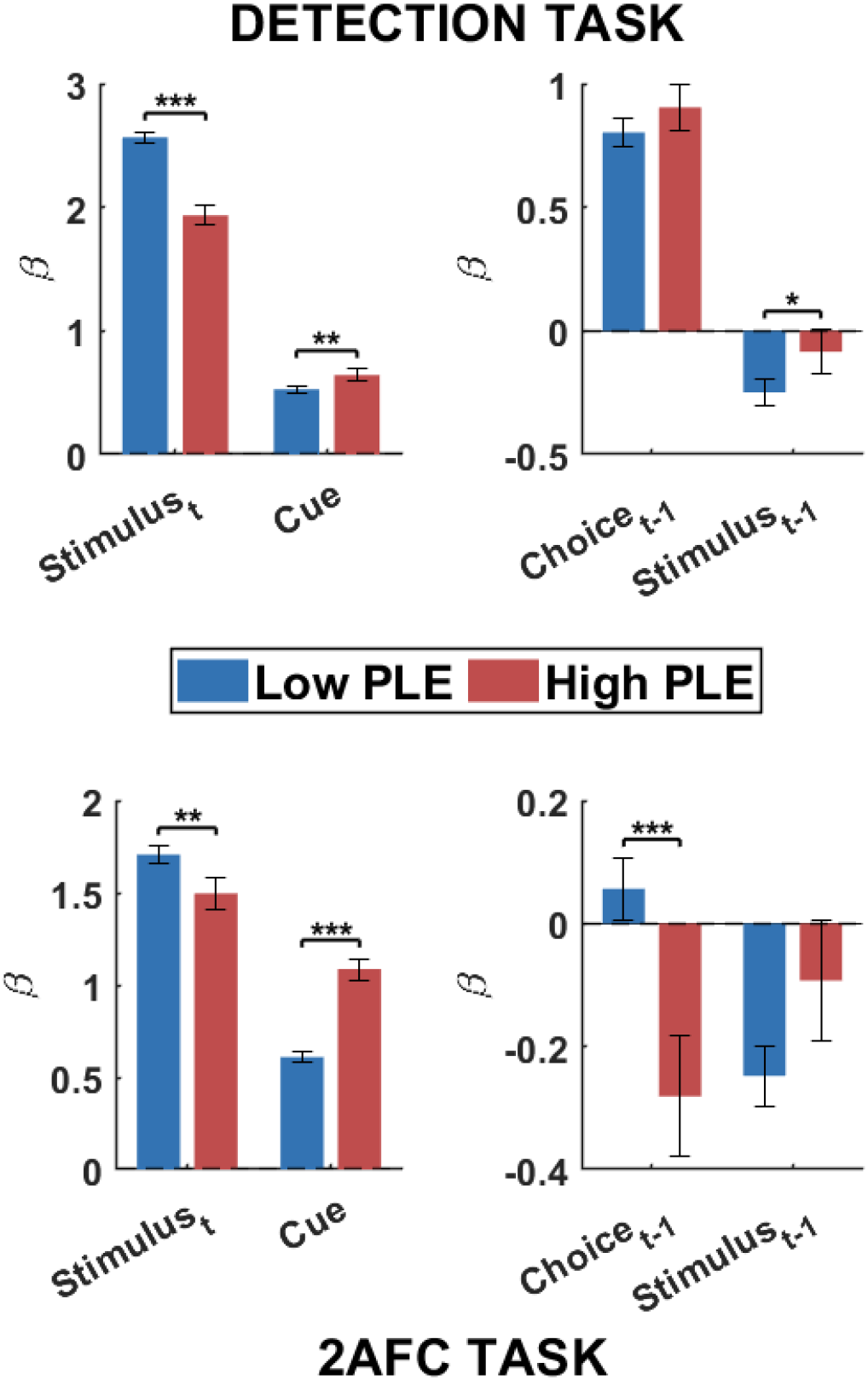
Modulation of perceptual biases by psychosis-like traits in two perceptual decision-making tasks. Bar plots (± SEM) illustrate how high-versus low-PLE participants differ in weighting sensory evidence, explicit cues, and choice history in the probabilistic detection task (top) and the random-dot motion discrimination task (bottom). In the detection task, those in the highest PLE tercile show a pronounced reduction in sensory evidence weighting (Stimulus_t_ × PLEgroup) yet maintain facilitatory cue use (Cue_t_ × PLEgroup). The response-repetition tendency remains unmodulated by PLE (Response_t−1_ × PLEgroup), while the repulsive carry-over from the previous stimulus is significantly attenuated in high-PLE group (Stimulus_t−1_ × PLEgroup). In the motion-discrimination task, high-PLE individuals again rely more heavily on cues (Cue_t_ × PLEgroup) while down-weighting sensory input (Stimulus_t_ × PLEgroup). Crucially, they reverse the response-repetition bias, exhibiting a significant shift away from their previous choice (Response_t−1_ × PLEgroup), whereas the repulsive stimulus-history effect shows only a non-significant trend (Stimulus_t−1_ × PLEgroup). Together, these results demonstrate that elevated psychosis-like traits consistently down-weight raw sensory evidence and up-weight prior expectations, but that their influence on sequential choice biases—attenuation versus inversion—varies according to task demands.

## Discussion

Our findings provide evidence that psychosis-like experiences (PLEs) systematically shape perceptual decision-making through specific alterations in sensory processing, predictive inference, and sequential decision biases. Using two complementary tasks—one probing detection under uncertainty and the other probing discrimination of motion direction—we show that elevated psychosis proneness is associated with reduced sensitivity to sensory evidence, heightened reliance on prior expectations, and a striking, context-specific modulation of choice history biases. These results converge with predictive processing theories of psychosis, which propose that disruptions in the hierarchical integration of priors and prediction errors underlie key clinical and subclinical phenomena (9,14,31).

Across both tasks, choice behaviour was positively influenced by current sensory input (Stimulus_t_) and cue probability (Cue_t_), indicating that perceptual decisions reflect the joint contribution of bottom-up evidence and top-down expectations. A consistent repulsive effect of the previous stimulus (Stimulus_t−1_) also emerged, suggesting a history-dependent mechanism that biases perception away from recent input. Notably, only in the detection task did the previous choice (Response_t−1_) significantly influence current decisions, revealing a tendency toward choice repetition that was absent in the motion discrimination task. These patterns suggest shared mechanisms for integrating evidence and prior information, alongside task-specific differences in response history effects. Crucially, the strength and direction of these influences were differentially modulated by PLE, affecting the balance between sensory evidence, prior information, and choice history.

First, consistent with a growing literature, we found that participants with higher PLE scores showed reduced influence of sensory input in shaping their decisions. This effect emerged robustly across both the detection and motion discrimination tasks and was further confirmed in tercile-based analyses. These findings resonate with prior work in schizophrenia and schizotypy indicating diminished sensory precision and hypo-weighting of bottom-up evidence during perceptual inference (12,14,28,32) but see (33). Our results extend these insights to subclinical populations, providing behavioral evidence that disruptions in the encoding or utilization of sensory information are not limited to clinical populations but are continuous across the psychosis spectrum (34). The reduction in sensory weighting may reflect aberrant computations in early sensory processing regions or altered functional connectivity with higher-order associative areas implicated in the representation of predictive priors (35–37). Indeed, previous neuroimaging research has documented reduced sensory cortical activation alongside increased top-down modulatory influence from frontal cortical regions in schizophrenia and schizotypy (35,38,39). Moreover, both schizophrenia and subclinical schizotypy have been associated with a lower individual alpha frequency (IAF) (40–42), a neural marker closely linked to the enhanced resolution of sensory evidence sampling (43,44). This suggests that the diminished reliance on sensory input observed in individuals with elevated PLEs may partly stem from disruptions in alpha oscillatory dynamics.

A complementary yet distinct finding was the enhanced reliance on predictive cues in participants with elevated psychosis-like traits. Across both experimental paradigms, psychosis-prone individuals consistently demonstrated exaggerated use of predictive priors to guide their perceptual decisions. Notably, the magnitude of the observed effect was robust across both continuous and categorical analyses of psychosis proneness, confirming the reliability and consistency of this computational alteration. This pattern further supports the predictive coding hypothesis, which suggests that psychosis involves an overly precise or rigid representation of internal expectations, overwhelming bottom-up sensory evidence (3,31). Our findings reinforce that a similar cognitive and computational substrate is already present subclinically, further validating psychosis-like traits as meaningful analogues of clinical psychosis.

A central contribution of this study is the demonstration that sequential decision biases—how recent perceptual history influence current ones—are not uniformly disrupted in psychosis proneness but instead flip direction depending on task demands. In the detection task, higher PLE scores were not associated with changes in response repetition but were linked to a marked attenuation of the repulsive bias from the previous stimulus—indicating a reduced influence of recent sensory history on behaviour. In contrast, in the motion discrimination task, high-PLE participants exhibited a significant repulsive bias relative to their previous choices, shifting away from their immediately preceding response, while the effect of the previous stimulus on current decisions remained unaffected by PLE scores. This striking double dissociation underscores that the influence of psychosis-proneness on choice history is not uniform, but varies as a function of the inferential demands imposed by different task structures.

This divergence cannot be explained by differences in task difficulty, as both tasks were calibrated to yield comparable accuracy (∼70%). Rather, it likely reflects differences in sensory and decision architecture between the two decisional contexts. Indeed, it is possible that the two tasks hinge on different levels of the perceptual hierarchy. In the detection task, repulsive effects from previous stimuli could be interpreted as reflecting low-level sensory adaptation—a form of neural gain control that suppresses responses to recently encountered features. Adaptation effect might be a consequence of one important goal of the visual system, i.e., to maximize sensitivity to changes in the physical environment (45). Our finding that this repulsion is selectively attenuated in high-PLE individuals suggests a deficit in early sensory mechanisms. This aligns with evidence of impaired adaptation and reduced gain control in schizophrenia-spectrum conditions (46,47) and suggests that subclinical traits may already manifest as reduced suppression of prior sensory traces. This interpretation could also fit with the highlighted robust negative interaction between stimulus in the current trial and PLE traits: individuals with elevated psychosis proneness tend to down-weight actual sensory input when forming a decision. Consequently, the sensory representation itself may be too degraded or noisy to elicit a reliable adaptive response, further dampening the typical repulsive after-effects associated with recent stimulation.

By contrast, in the motion discrimination task—where perceptual decisions depend on the accumulation of noisy evidence over time—a different form of bias emerges. High-PLE participants exhibit a significant repulsive bias from their own previous response, tending to shift away from their last choice. In contrast, low-PLE individuals appear to show the opposite tendency, displaying an attractive bias toward repeating the previous response.

One possible interpretation is that psychosis-prone individuals adopt a compensatory strategy at the decisional level, actively avoiding response repetition. This pattern is consistent with previous reports of increased belief instability (48) and an overestimation of context change in individuals with schizophrenia (49). Such an overestimation may lead psychosis-prone individuals to infer that the correct response on the next trial is likely to differ from the previous one. Crucially, this double dissociation—reduced stimulus-driven repulsion in the detection task versus decision-level repulsion in the motion task—mirrors prior evidence for two distinct history‐bias mechanisms: a fast, repulsive sensory adaptation mechanism and a slower, attractive bias toward previous trials, reflecting decision-level stability (45). Our findings suggest that psychosis proneness not only dampens the former but appears to invert the latter—transforming what is typically an attractive decisional bias into a repulsive one.

Moreover, Fritsche et al. (45) demonstrated that the attractive decision bias likely originates from a post-perceptual shift in working memory representations, whereby the current stimulus is biased toward previous perceptual decisions. This is consistent with earlier reports of trial-to-trial carryover effects in short-term memory (50). The inversion observed in high-PLE individuals may also reflect a deficit in maintaining stable representations of prior decisions, possibly due to reduced working memory (WM) capacity (51), which in turn would lead to decreased reliance on information from the previous trial. Crucially, Xie et al., (52) showed that individuals with more schizotypal features retained less precise representations in visual WM without a significant reduction in the number of retained WM representations.

An alternative account is that the observed sequential biases reflect a reliance on simple heuristics to guide decision-making. High-PLE individuals might engage in a form of “lose-shift” behaviour, wherein the tendency to switch response reflects an internal inference that the previous decision was likely incorrect, despite the absence of explicit feedback. Such behaviour presupposes the existence of implicit confidence signals that inform subsequent action based on internal estimates of uncertainty. While this strategy can be adaptive in volatile environments, it may become maladaptive when based on an unstable or miscalibrated internal model of environmental states (31). At first glance, this interpretation seems at odds with reports of overconfidence in schizophrenia (53) and positive schizotypy (54), which would instead predict a stronger tendency to repeat past choices —a “win-stay” strategy. However, such overconfidence may reflect a distorted metacognitive judgment acting at an explicit level, whereas lower-level perceptual confidence signals may be impaired. This dissociation could undermine the stability of internal decision representations, thereby promoting repulsive choice history effects. Consistent with this view, animal studies have shown that confidence-related signals originate in the same neural populations involved in evidence accumulation during perceptual decision-making, such as neurons in the lateral intraparietal area (LIP) during random dot motion tasks (55,56). Moreover, the human homolog of this region—the intraparietal sulcus—has also been causally implicated in metacognitive processes (57). This suggests that, in psychosis-prone individuals, implicit confidence signals arising during evidence accumulation may fail to reliably reflect the quality of incoming sensory information. As a result, decisions may be updated in an overly reactive or unstable manner, leading to a greater tendency to shift away from previous choices.

While this study offers important insights, several considerations merit acknowledgment. First, although we examined psychosis-proneness dimensionally and categorically, future research could benefit from longitudinal designs to understand whether observed computational alterations predict transition from subclinical psychosis-like experiences to clinical psychosis symptoms. Additionally, integrating neuroimaging methods could further clarify the neural substrates underlying reduced sensory precision and enhanced prior reliance observed here, particularly regarding connectivity between early sensory and high-level predictive processing regions.

Furthermore, examining the generalizability of our findings across sensory modalities and cognitive domains remains a critical direction for future research. While the present results suggest robust effects of psychosis-like traits on visual decision-making, recent evidence indicates that these effects may not extend uniformly across perceptual systems. For instance, Eckert et al. (20) found that, despite participants with higher psychosis proneness showing increased reliance on probabilistic priors in a 2AFC visual task—akin to our motion discrimination paradigm—no such effect emerged when the same probabilistic structure was applied to auditory stimuli. This sensory-specific discrepancy highlights the need to better understand how predictive processing disruptions may differentially manifest across perceptual modalities. It also raises the possibility that the weighting of priors and sensory evidence in psychosis is shaped not only by trait-like cognitive styles but also by modality-specific constraints or differences in the neural architecture supporting perception in different sensory systems.

Finally, by revealing dissociable alterations in how sensory evidence, prior information, and choice history are integrated across contexts, our results point to potential targets for cognitive or neuromodulatory interventions. Approaches aimed at restoring the balance between external evidence and internal expectations—such as recalibrating sensory precision or modulating the weighting of prior beliefs—may hold promise in mitigating early perceptual disturbances along the psychosis continuum.

In conclusion, our study provides robust and comprehensive evidence that psychosis-like traits significantly modulate perceptual decision-making processes, characterized by reduced sensory precision, increased reliance on prior expectations, and context-dependent alterations in history biases. These findings substantiate predictive processing models of psychosis, highlighting nuanced computational mechanisms underlying psychosis proneness, with significant theoretical and clinical implications. By revealing dissociable and task-dependent disruptions in predictive inference, our results highlight the need for a nuanced, multidimensional framework to understand how psychosis vulnerability manifests across different cognitive domains—both in clinical populations and along the broader psychosis continuum.

## References

1. Clark A (2013): Whatever next? Predictive brains, situated agents, and the future of cognitive science. Behav Brain Sci 36: 181–204.

2. Friston K, Kiebel S (2009): Predictive coding under the free-energy principle. Philos Trans R Soc Lond B Biol Sci 364: 1211– 1221.

3. Powers AR, Mathys C, Corlett PR (2017): Pavlovian conditioning-induced hallucinations result from overweighting of perceptual priors. Science 357: 596–600.

4. Schmack K, Castro AG-C de, Rothkirch M, Sekutowicz M, Rössler H, Haynes J-D, et al. (2013): Delusions and the Role of Beliefs in Perceptual Inference. J Neurosci 33: 13701–13712.

5. Adams RA, Stephan KE, Brown HR, Frith CD, Friston KJ (2013): The computational anatomy of psychosis. Front Psychiatry 4: 47.

6. Tarasi L, Trajkovic J, Diciotti S, di Pellegrino G, Ferri F, Ursino M, Romei V (2022): Predictive waves in the autism-schizophrenia continuum: A novel biobehavioral model. Neurosci Biobehav Rev 132: 1–22.

7. Kafadar E, Fisher VL, Quagan B, Hammer A, Jaeger H, Mourgues C, et al. (2022): Conditioned Hallucinations and Prior Overweighting Are State-Sensitive Markers of Hallucination Susceptibility. Biol Psychiatry 92: 772–780.

8. Corlett PR, Frith CD, Fletcher PC (2009): From drugs to deprivation: a Bayesian framework for understanding models of psychosis. Psychopharmacology 206: 515–530.

9. Corlett PR, Horga G, Fletcher PC, Alderson-Day B, Schmack K, Powers AR (2019): Hallucinations and Strong Priors. Trends Cogn Sci 23: 114–127.

10. Nelson B, Yuen HP, Wood SJ, Lin A, Spiliotacopoulos D, Bruxner A, et al. (2013): Long-term follow-up of a group at ultra high risk (“prodromal”) for psychosis: the PACE 400 study. JAMA Psychiatry 70: 793–802.

11. van Os J, Reininghaus U (2016): Psychosis as a transdiagnostic and extended phenotype in the general population. World Psychiatry 15: 118–124.

12. Benrimoh D, Fisher VL, Seabury R, Sibarium E, Mourgues C, Chen D, Powers A (2024): Evidence for Reduced Sensory Precision and Increased Reliance on Priors in Hallucination-Prone Individuals in a General Population Sample. Schizophrenia Bulletin 50: 349–362.

13. Stuke H, Kress E, Weilnhammer VA, Sterzer P, Schmack K (2021): Overly Strong Priors for Socially Meaningful Visual Signals Are Linked to Psychosis Proneness in Healthy Individuals. Front Psychol 12. 10.3389/fpsyg.2021.583637

14. Tarasi L, Martelli ME, Bortoletto M, di Pellegrino G, Romei V (2023): Neural Signatures of Predictive Strategies Track Individuals Along the Autism-Schizophrenia Continuum. Schizophr Bull 49: 1294–1304.

15. Teufel C, Subramaniam N, Dobler V, Perez J, Finnemann J, Mehta PR, et al. (2015): Shift toward prior knowledge confers a perceptual advantage in early psychosis and psychosis-prone healthy individuals. Proc Natl Acad Sci U S A 112: 13401–13406.

16. Cicchini GM, Mikellidou K, Burr DC (2024): Serial Dependence in Perception. Annual Review of Psychology 75: 129–154.

17. Pascucci D, Tanrikulu ÖD, Ozkirli A, Houborg C, Ceylan G, Zerr P, et al. (2023): Serial dependence in visual perception: A review. Journal of Vision 23: 9.

18. Ranieri G, Benedetto A, Ho HT, Burr DC, Morrone MC (2022): Evidence of Serial Dependence from Decoding of Visual Evoked Potentials. J Neurosci 42: 8817–8825.

19. Urai AE, de Gee JW, Tsetsos K, Donner TH (2019): Choice history biases subsequent evidence accumulation ((T. Verstynen, B. G. Shinn-Cunningham, & T. Verstynen, editors)). eLife 8: e46331.

20. Eckert A-L, Gounitski Y, Guggenmos M, Sterzer P (2023): Cross-Modality Evidence for Reduced Choice History Biases in Psychosis-Prone Individuals. Schizophrenia Bulletin 49: 397–406.

21. Stein H, Barbosa J, Rosa-Justicia M, Prades L, Morató A, Galan-Gadea A, et al. (2020): Reduced serial dependence suggests deficits in synaptic potentiation in anti-NMDAR encephalitis and schizophrenia. Nat Commun 11: 4250.

22. Streiling K, Schülke R, Straube B, van Dam LCJ (2025): Choice- and trial-history effects on causality perception in Schizophrenia Spectrum Disorder. Schizophr 11: 1–11.

23. Pascucci D, Roinishvili M, Chkonia E, Brand A, Whitney D, Herzog MH, Manassi M (2025): Intact Serial Dependence in Schizophrenia: Evidence from an Orientation Adjustment Task. Schizophrenia Bulletin 51: 754–764.

24. Gold JI, Shadlen MN (2007): The neural basis of decision making. Annu Rev Neurosci 30: 535–574.

25. Tarasi L, di Pellegrino G, Romei V (2022): Are you an empiricist or a believer? Neural signatures of predictive strategies in humans. Progress in Neurobiology 219: 102367.

26. Tarasi L, Bertaccini R, Ippolito G, Martelli ME, di Pellegrino G, Romei V (2025): Oscillatory signatures of monitoring and anticipatory strategies for probabilistic vs deterministic cues. Imaging Neuroscience 3: imag_a_00496.

27. Tarasi L, Romei V (2024): Individual Alpha Frequency Contributes to the Precision of Human Visual Processing. Journal of Cognitive Neuroscience 36: 602–613.

28. Tarasi L, De Fatis CT, Covelli M, Ippolito G, Avenanti A, Romei V (2025): Preparing to act follows Bayesian inference rules. iScience. 10.1101/2024.08.16.608232

29. Wichmann FA, Hill NJ (2001): The psychometric function: I. Fitting, sampling, and goodness of fit. Perception & Psychophysics 63: 1293–1313.

30. Raine A (1991): The SPQ: A Scale for the Assessment of Schizotypal Personality Based on DSM-III-R Criteria. Schizophrenia Bulletin 17: 555–564.

31. Sterzer P, Adams RA, Fletcher P, Frith C, Lawrie SM, Muckli L, et al. (2018): The Predictive Coding Account of Psychosis. Biol Psychiatry 84: 634–643.

32. Benrimoh D, Parr T, Vincent P, Adams RA, Friston K (2018): Active Inference and Auditory Hallucinations. Comput Psychiatr 2: 183–204.

33. Schaub A-C, Eckert A-L, Nuiten S, Weilnhammer V, Sterzer P (2025, March 24): Reduced weighting of short-term perceptual priors during auditory perceptual decision-making in psychosis-prone individuals. OSF. 10.31234/osf.io/7zgqk_v2

34. van Os J, Reininghaus U (2016): Psychosis as a transdiagnostic and extended phenotype in the general population. World Psychiatry 15: 118–124.

35. Alamia A, Gordillo D, Chkonia E, Roinishvili M, Cappe C, Herzog MH (2024): Oscillatory traveling waves provide evidence for predictive coding abnormalities in schizophrenia. Biological Psychiatry. 10.1016/j.biopsych.2024.11.014

36. Tarasi L, Alamia A, Romei V (2025, January 16): Backward alpha oscillations shape perceptual bias under probabilistic cues. bioRxiv, p 2025.01.14.632925.

37. Ippolito G, Bertaccini R, Tarasi L, Di Gregorio F, Trajkovic J, Battaglia S, Romei V (2022): The Role of Alpha Oscillations among the Main Neuropsychiatric Disorders in the Adult and Developing Human Brain: Evidence from the Last 10 Years of Research [no. 12]. Biomedicines 10: 3189.

38. Schmack K, Castro AG-C de, Rothkirch M, Sekutowicz M, Rössler H, Haynes J-D, et al. (2013): Delusions and the Role of Beliefs in Perceptual Inference. J Neurosci 33: 13701–13712.

39. Schmack K, Rothkirch M, Priller J, Sterzer P (2017): Enhanced predictive signalling in schizophrenia. Human Brain Mapping 38: 1767–1779.

40. Ramsay IS, Lynn PA, Schermitzler B, Sponheim SR (2021): Individual alpha peak frequency is slower in schizophrenia and related to deficits in visual perception and cognition. Sci Rep 11: 17852.

41. Tarasi L, Romanazzi D, Pasini A, Romei V (2025): Delusion-like thinking is associated with lower individual alpha peak frequency. Schizophrenia.

42. Trajkovic J, Di Gregorio F, Ferri F, Marzi C, Diciotti S, Romei V (2021): Resting state alpha oscillatory activity is a valid and reliable marker of schizotypy. Sci Rep 11: 10379.

43. Di Gregorio F, Trajkovic J, Roperti C, Marcantoni E, Di Luzio P, Avenanti A, et al. (2022): Tuning alpha rhythms to shape conscious visual perception. Current Biology 32: 988-998.e6.

44. Samaha J, Postle BR (2015): The Speed of Alpha-Band Oscillations Predicts the Temporal Resolution of Visual Perception. Current Biology 25: 2985–2990.

45. Fritsche M, Mostert P, de Lange FP (2017): Opposite Effects of Recent History on Perception and Decision. Curr Biol 27: 590–595.

46. Cornelis C, De Picker LJ, Coppens V, Morsel A, Timmers M, Dumont G, et al. (2021): Impaired Sensorimotor Adaption in Schizophrenia in Comparison to Age-Matched and Elderly Controls. Neuropsychobiology 81: 127–140.

47. Thakkar KN, Silverstein SM, Brascamp JW (2019): A review of visual aftereffects in schizophrenia. Neuroscience & Biobehavioral Reviews 101: 68–77.

48. Hauke DJ, Roth V, Karvelis P, Adams RA, Moritz S, Borgwardt S, et al. (2022): Increased Belief Instability in Psychotic Disorders Predicts Treatment Response to Metacognitive Training. Schizophrenia Bulletin 48: 826–838.

49. Kaplan CM, Saha D, Molina JL, Hockeimer WD, Postell EM, Apud JA, et al. (2016): Estimating changing contexts in schizophrenia. Brain 139: 2082–2095.

50. Visscher KM, Kahana MJ, Sekuler R (2009): Trial-to-trial carryover in auditory short-term memory. Journal of Experimental Psychology: Learning, Memory, and Cognition 35: 46–56.

51. Farmer CM, O’Donnell BF, Niznikiewicz MA, Voglmaier MM, McCarley RW, Shenton ME (2000): Visual Perception and Working Memory in Schizotypal Personality Disorder. AJP 157: 781–788.

52. Xie W, Cappiello M, Park H-B, Deldin P, Chan RCK, Zhang W (2018): Schizotypy is associated with reduced mnemonic precision in visual working memory. Schizophrenia Research 193: 91–97.

53. Moritz S, Ramdani N, Klass H, Andreou C, Jungclaussen D, Eifler S, et al. (2014): Overconfidence in incorrect perceptual judgments in patients with schizophrenia. Schizophrenia Research: Cognition 1: 165–170.

54. Lehmann M, Ettinger U (2023): Metacognitive monitoring in schizotypy: Systematic literature review and new empirical data. J Behav Ther Exp Psychiatry 81: 101891.

55. Kiani R, Shadlen MN (2009): Representation of confidence associated with a decision by neurons in the parietal cortex. Science 324: 759–764.

56. Zylberberg A, Shadlen MN (2025): A population representation of the confidence in a decision in the parietal cortex. Cell Rep 44: 115526.

57. Luzio PD, Tarasi L, Silvanto J, Avenanti A, Romei V (2022): Human perceptual and metacognitive decision-making rely on distinct brain networks. PLOS Biology 20: e3001750.

